# An archaeal symbiont-host association from the deep terrestrial subsurface

**DOI:** 10.1101/486977

**Authors:** Katrin Schwank, Till L. V. Bornemann, Nina Dombrowski, Anja Spang, Jillian F. Banfield, Alexander J. Probst

## Abstract

DPANN archaea have reduced metabolic capacities and are diverse and abundant in deep aquifer ecosystems, yet little is known about their interactions with other microorganisms that reside there. Here, we provide evidence for an archaeal host-symbiont association from a deep aquifer system at the Colorado Plateau (Utah, USA). The symbiont, Candidatus *Huberiarchaeum crystalense*, and its host, Ca. *Altiarchaeum hamiconexum*, show a highly significant co-occurrence pattern over 65 metagenome samples collected over six years. The physical association of the two organisms was confirmed with genome-informed fluorescence *in situ* hybridization depicting small cocci of Ca. *H. crystalense* attached to Ca. *A. hamiconexum* cells. Based on genomic information, Ca. *H. crystalense* has a similar metabolism as *Nanoarchaeum equitans* and potentially scavenges vitamins, sugars, nucleotides, and reduced redox-equivalents from its host. These results provide insight into host-symbiont interactions among members of two uncultivated archaeal phyla that thrive in a deep subsurface aquifer.

---

The DPANN (Diapherotrites, Parvarchaeota, Aigarchaeota, Nanoarchaeota, Nanohaloarchaeota) [1] radiation is a proposed monophyletic group of diverse archaeal phyla whose organisms exhibit mainly reduced genomes with limited metabolic capacities [2]. While most of these archaea were suggested to live in symbiosis with other microorganisms, respective hosts were only described for Nanoarchaeota [3, 4] and Micrarchaeota (ARMAN) symbionts [5–7]. However, no DPANN-host interaction has been described for representatives in aquifer systems, where these archaea are particularly abundant and very diverse [2].

Recently, subsurface fluids discharged by a cold, CO_2_-driven geyser at the Colorado Plateau, Utah (Crystal Geyser) revealed the presence of multiple novel archaeal organisms, including the first representative of the Candidatus phylum Huberarchaeota [8], which we show to be part of the DPANN radiation (*Fig. 1A*, *Supplementary Data Files 1-5*). Interestingly, the only representative, Ca. *Huberiarchaeum crystalense*, has previously been shown to correlate in relative genome abundance with the most abundant primary producer in the ecosystem, Ca. *Altiarchaeum hamiconexum* CG over five days [8]. To validate this initial inference, we leveraged 65 metagenomic samples taken between 2009 and 2015. This not only confirmed the correlation (*Fig. 1B*, Supplementary Methods) but also provided evidence that both archaea occur mainly on filters with large pore sizes (>0.65 μm), although the *Altiarchaeum* cell sizes can be as small as 0.4 μm [9]. Consequently, we tested the hypothesis that both species appeared as cell aggregates in groundwater. We investigated groundwater samples collected onto a 0.2 μm filter taken during geyser eruptions when the putative host *A. hamiconexum* CG showed the greatest abundance based on genome read-coverage. We designed an oligonucleotide probe for the 16S rRNA gene of Ca. *H. crystalense* and evaluated it against other 16S rRNA genes reconstructed from metagenomes [8] (see Supplementary Methods). The application of this specific probe in MiL-FISH (multilabeled fluorescence *in-situ* hybridization [10]; *Fig. 1C*) along with a general probe for Altiarchaeota [11] showed not only the appearance of individual cells for each archaeon (*Fig. 1C-I/II*), but also the co-localization of the two archaea (*Fig. 1C-III/IV*). The putative symbiont Ca. *H. crystalense* was identified to occur as an episymbiont attached to Ca. *A. hamiconexum* CG (*Fig. 1C-III*), sometimes forming larger cell aggregates with the host (*Fig. 1C-IV*) explaining their occurrence mainly on filters with larger pore sizes. The median ratio of Altiarchaeota and Huberarchaeota genome abundances across the different samples was about 11:1 in bulk samples, indicating that the host-symbiont relationship is not only constant but also that not all Altiarchaeota carry a symbiont (*Fig. S1*). Although we did detect more individual Altiarchaeota than consortia with Huberarchaeota in FISH analyses, the ratio could not be confirmed statistically due to the extremely low cell numbers in the groundwater and the heavy deposits of minerals on the filters.

**Figure 1.**
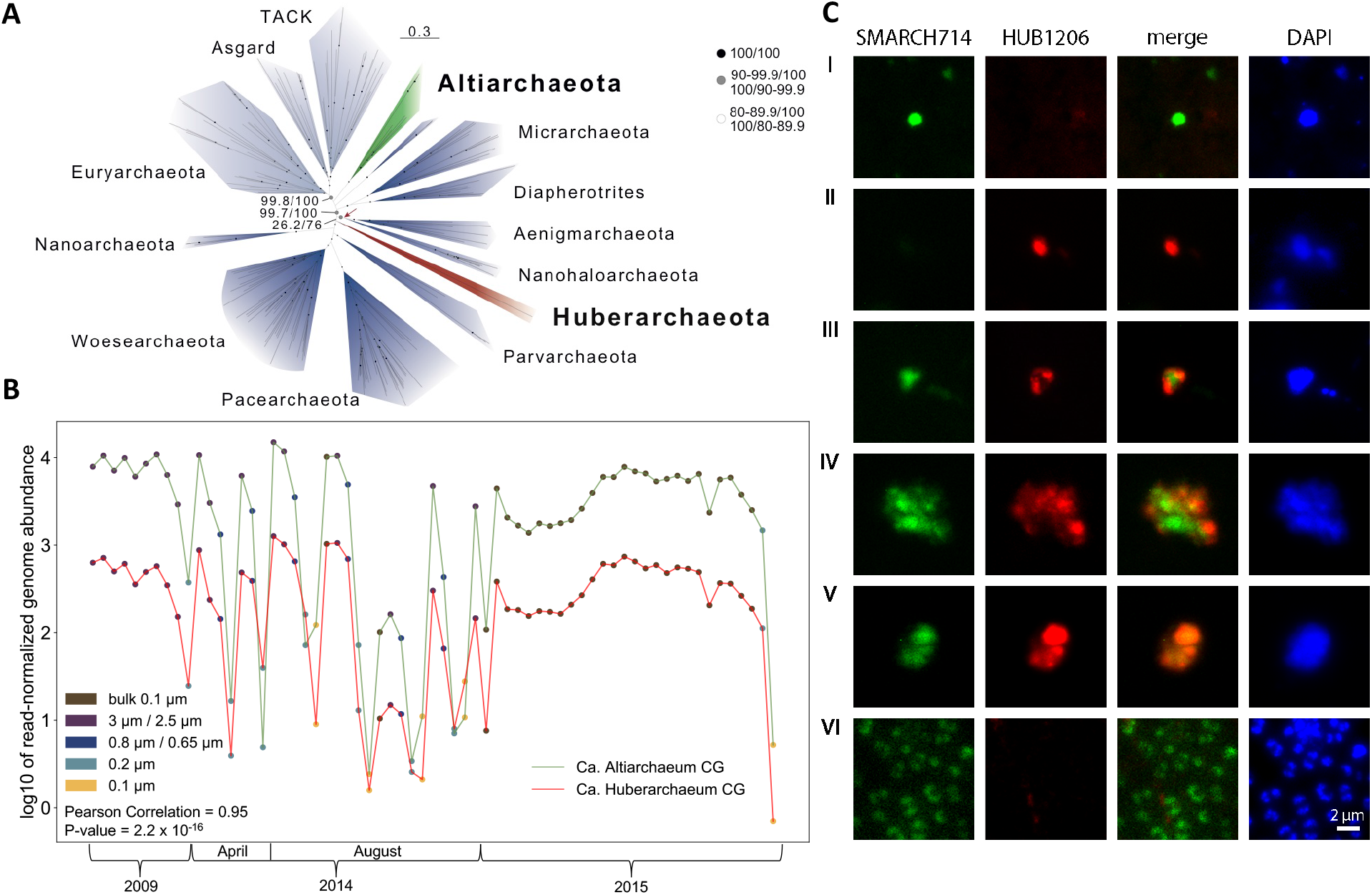
Phylogeny and co-occurrence analysis of Ca. *Huberiarchaeum crystalense* and Ca. *Altiarchaeum hamiconexum* CG. A. Phylogenetic placement of Huberarchaeota using a concatenated protein alignment of 34 markergenes. This maximum likelihood tree includes 186 taxa and was inferred based on an alignment of 4224 positions in IQ-tree under the LG+C60+F+R model. The green arrow highlights the alternative placement of the Huberarchaeota in the IQ-tree analyses based on an SR4 recoded alignment (support: 6.2/76). Bootstrap support was inferred using a SH-like approximate likelihood ratio test and ultrafast bootstrap support values and are represented by small circles according to the legend. Scale bar indicates the average number of substitutions per site. For details on the methods please see supplementary data. B. Co-correlation analysis of relative abundance of genomes of Ca. *Huberiarchaeum crystalense* and *Altiarchaeum hamiconexum* CG based on 65 metagenome samples taken between 2009 and 2015. Interestingly, the co-occurrence also holds true between samples taken with filters of different pore-sizes indicating that the organisms form an association that does not pass through small pore sizes. C. MiL fluorescence *in situ* hybridization depicting the association of Ca. *Huberiarchaeum crystalense* and *Altiarchaeum hamiconexum* CG cells and the respective controls. SMARCH714 is a specific probe for Altiarchaeota [11] labeled with Atto488. HUB1206 is a probe designed off the 16S rRNA gene sequence of Huberiarchaeum based on information from multiple genomic bins (see *Supplementary Methods*). The probe was labeled with CY3. **I**. Individual *Altiarchaeum hamiconexum* CG cell. **II**. Individual Ca. *H. crystalense* cell. **III**. Attachment of Ca. *H. crystalense* to *Altiarchaeum hamiconexum* CG **IV**. occurrence of Ca. *H. crystalense* to *Altiarchaeum hamiconexum* CG in cell clusters **V**. Overlapping fluorescence signals of the two probes putatively in the same cells. **IV**. Control sample of a well-studied ecosystem dominated by *Altiarchaeum hamiconexum* IMS [19], where according to metagenomic data no Huberarchaeota occur. We show that the specific probe of Ca. *Huberiarchaeum crystalense* does not bind to Altiarchaeota or other organisms in the ecosystem.

To further investigate the potential metabolic interactions as well as the dependency of Ca. *H. crystalense* on Ca. *A. hamiconexum* CG, we analyzed [12] the metabolic capacity encoded by their respective pangenomes *in silico* (see *Supplementary Methods; Fig. 2*). Metabolic predictions based on the Ca. *H. crystalense* pangenome indicate that this organisms heavily relies on diverse metabolic intermediates from its host and the predicted metabolic pathways showed great similarity to those found for *Nanoarchaeum equitans* [13]. For instance, Ca. *H. crystalense* cannot conserve energy in the form of ATP or GTP through oxidative phosphorylation, as its genomes do not encode any subunit of the ATP-Synthase complex. Similarly, we did not detect any genes involved in substrate level phosphorylation. Although the *de novo* synthesis of amino acids is incomplete in Ca. *H. crystalense*, its genome encodes for six proteases and peptidases (Table S1), which could break down proteins and retrieve necessary amino acids. Pathways for converting aromatic amino acids into one another exist in its genome and potentially enable some metabolic flexibility. While transcription and DNA replication (including DNA repair) are possible in Ca. *H. crystalense*, required nucleotides for these processes cannot be synthesized *de novo* and must thus be acquired from the environment, likely from its host. Similar to *N. equitans* [13], Ca. *H. crystalense* encodes for very few recognizable transport systems. Activated sugars and vitamins cannot be synthesized either, yet are essential for multiple enzymes and anabolic processes (e.g., glycosyltransferases and aminotransferases).

**Figure 2.**
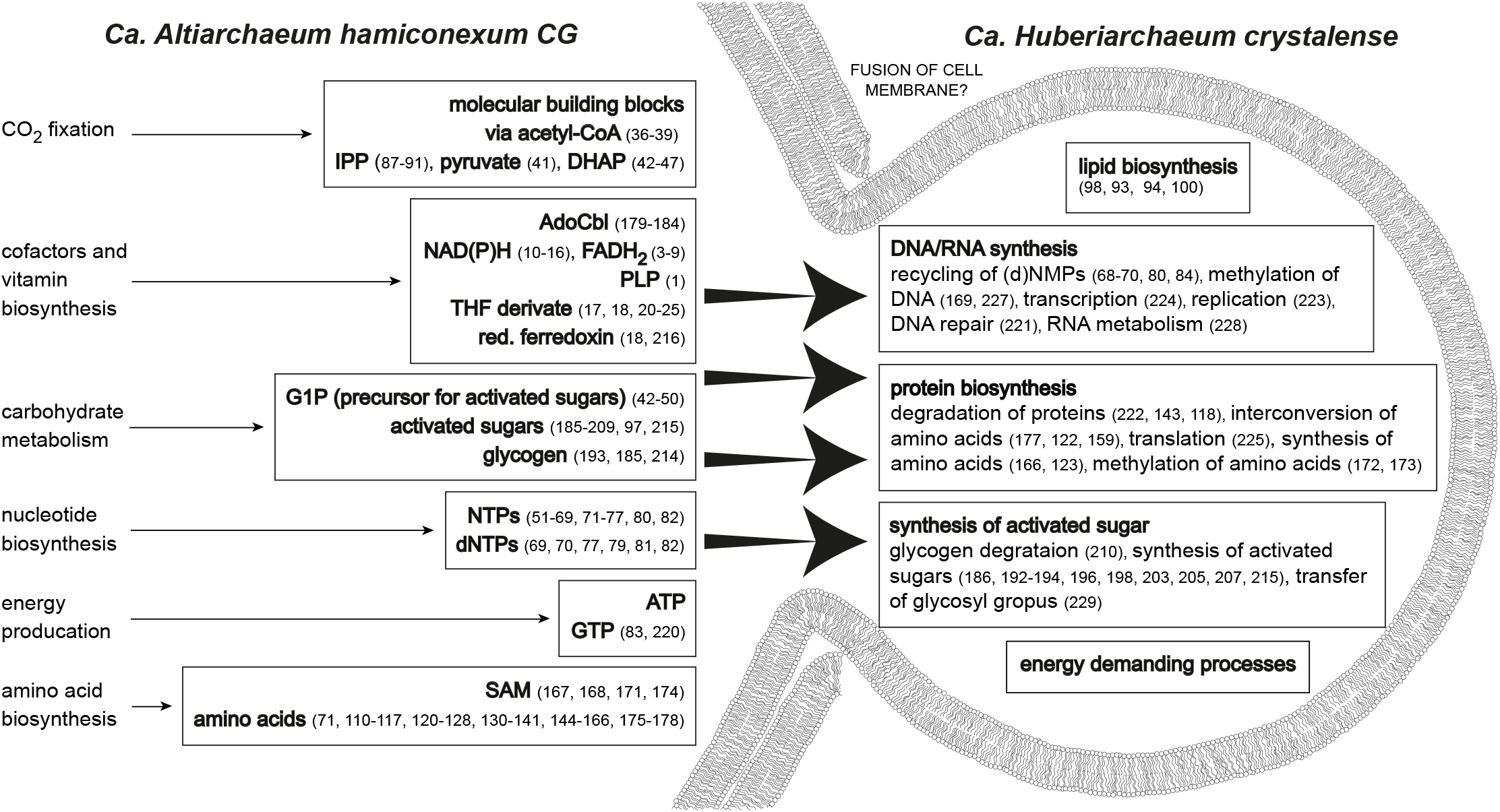
Potential metabolic interactions of Ca. *Altiarchaeum hamiconexum* CG and Ca. *Huberiarchaeum crystelense*. The depicted figure is based on a putative fusion of the membranes as it was reported for *N. equitans* and *I. hospitalis* [20]. Numbers refer to predicted enzymes involved in the respective metabolic function (Supplementary Table 1 and 2). Individual, predicted enzymatic reactions are depicted in Fig. S1.

Ca. *H. crystalense* does not have the capacity to synthesize redox-equivalents like NAD(P)^+^ or ferredoxin, although both are required by multiple enzymes encoded in its genome. Importantly, while we did not detect the capacity to reduce ferredoxin, some encoded enzymes necessitate this iron sulfur protein as a co-factor. For instance, digernaylgeranylglycophospholipid reductase was previously shown to only operate with reduced ferredoxin in archaea [14]. This enzyme participates in archaeal lipid biosynthesis, a nearly complete pathway in Ca. *H. crystalense*. This is in stark contrast to *N. equitans*, which is incapable of lipid metabolism and requires lipids from its host [13] and also differs from other members of the DPANN, most of which lack genes encoding lipid biosynthesis proteins [15]. Correlations of lipid and metagenome analyses suggested that Ca. *H. crystalense* has a very specific lipid different from its host [16], supporting the notion that lipid biosynthesis is indeed carried out by this archaeon. Hence, the cells must obtain ferredoxin in a reduced state. However, reduced ferredoxin (Eo’^~^-500mV) cannot freely exist in deep groundwater fluids of Crystal Geyser (pH = 6.8, T= 17°C). Thus, reduced ferredoxin might be retrieved from the host for example via vesicles or cell fusions. Some FISH images showed an overlap of signals from Ca. *H. crystalense* and Ca. *A. hamiconexum* CG, suggesting the possibility of direct cytoplasmic contact (*Fig. 1C-V*). For instance, surface proteins homologous to hemolysine [8] could initiate such contacts as observed previously for acidophilic Micrarchaeota using cryo-electron tomography [5, 17]. More recently, fusion of cell membranes of *N. equitans* and its host *Ignicoccus hospitalis* was observed by transmission electron microscopy [18].

Altogether, this first discovery of an archaeal symbiont-host association from an aquifer system provides insights into metabolic capacities and interactions of a DPANN archaeon together with its host in the deep terrestrial subsurface. The host’s metabolism appeared to complement the symbiont’s needs for diverse metabolic intermediates. Apart from the lipid metabolism, the metabolic capacities of Ca. *H. crystalense* are similar to those of *N. equitans* indicating convergent evolutionary processes in both symbiont-host associations.

## ACKNOWLEDGEMENTS

This study was funded by the Ministerium für Kultur und Wissenschaft des Landes Nordrhein-Westfalen (“Nachwuchsgruppe Dr. Alexander Probst”). A.S. is supported by a grant of the Swedish Research Council (VR starting grant 2016-) and the NWO-I foundation of the Netherlands Organisation for Scientific Research (WISE fellowship).

## References

1. Rinke C, Schwientek P, Sczyrba A, Ivanova NN, Anderson IJ, Cheng J-F, et al. Insights into the phylogeny and coding potential of microbial dark matter. Nature 2013; 499: 431–437.

2. Castelle CJ, Wrighton KC, Thomas BC, Hug LA, Brown CT, Wilkins MJ, et al. Genomic expansion of domain archaea highlights roles for organisms from new phyla in anaerobic carbon cycling. Curr Biol CB 2015; 25: 690–701.

3. Huber H, Hohn MJ, Rachel R, Fuchs T, Wimmer VC, Stetter KO. A new phylum of Archaea represented by a nanosized hyperthermophilic symbiont. Nature 2002; 417: 63–67.

4. Jarett JK, Nayfach S, Podar M, Inskeep W, Ivanova NN, Munson-McGee J, et al. Single-cell genomics of co-sorted Nanoarchaeota suggests novel putative host associations and diversification of proteins involved in symbiosis. Microbiome 2018; 6: 161.

5. Baker BJ, Comolli LR, Dick GJ, Hauser LJ, Hyatt D, Dill BD, et al. Enigmatic, ultrasmall, uncultivated Archaea. Proc Natl Acad Sci 2010; 200914470.

6. Krause S, Bremges A, Münch PC, McHardy AC, Gescher J. Characterisation of a stable laboratory co-culture of acidophilic nanoorganisms. Sci Rep 2017; 7: 3289.

7. Golyshina OV, Toshchakov SV, Makarova KS, Gavrilov SN, Korzhenkov AA, La Cono V, et al. ‘ARMAN’archaea depend on association with euryarchaeal host in culture and in situ. Nat Commun 2017; 8: 60.

8. Probst AJ, Ladd B, Jarett JK, Geller-McGrath DE, Sieber CMK, Emerson JB, et al. Differential depth distribution of microbial function and putative symbionts through sediment-hosted aquifers in the deep terrestrial subsurface. Nat Microbiol 2018; 3: 328–336.

9. Moissl C, Rudolph C, Rachel R, Koch M, Huber R. In situ growth of the novel SM1 euryarchaeon from a string-of-pearls-like microbial community in its cold biotope, its physical separation and insights into its structure and physiology. Arch Microbiol 2003; 180: 211–217.

10. Schimak MP, Kleiner M, Wetzel S, Liebeke M, Dubilier N, Fuchs BM. MiL-FISH: Multilabeled oligonucleotides for fluorescence in situ hybridization improve visualization of bacterial cells. Appl Env Microbiol 2016; 82: 62–70.

11. Rudolph C, Wanner G, Huber R. Natural communities of novel archaea and bacteria growing in cold sulfurous springs with a string-of-pearls-like morphology. Appl Environ Microbiol 2001; 67: 2336–2344.

12. Vallenet D, Labarre L, Rouy Z, Barbe V, Bocs S, Cruveiller S, et al. MaGe: a microbial genome annotation system supported by synteny results. Nucleic Acids Res 2006; 34: 53–65.

13. Waters E, Hohn MJ, Ahel I, Graham DE, Adams MD, Barnstead M, et al. The genome of Nanoarchaeum equitans: Insights into early archaeal evolution and derived parasitism. Proc Natl Acad Sci 2003; 100: 12984–12988.

14. Isobe K, Ogawa T, Hirose K, Yokoi T, Yoshimura T, Hemmi H. Geranylgeranyl reductase and ferredoxin from Methanosarcina acetivorans are required for the synthesis of fully reduced archaeal membrane lipid in Escherichia coli cells. J Bacteriol 2014; 196: 417–423.

15. Villanueva L, Schouten S, Damsté JSS. Phylogenomic analysis of lipid biosynthetic genes of Archaea shed light on the ‘lipid divide’. Environ Microbiol 2017; 19: 54–69.

16. Probst AJ, Elling FJ, Castelle CJ, Zhu Q, Elvert M, Birarda G, et al. Lipid analysis of CO_2_-rich subsurface aquifers suggests an autotrophy-based deep biosphere with lysolipids enriched in CPR bacteria. bioRxiv 2018; 465690.

17. Comolli LR, Baker BJ, Downing KH, Siegerist CE, Banfield JF. Three-dimensional analysis of the structure and ecology of a novel, ultra-small archaeon. ISME J 2009; 3: 159.

18. Heimerl T, Flechsler J, Pickl C, Heinz V, Salecker B, Zweck J, et al. A complex endomembrane system in the archaeon *Ignicoccus hospitalis* tapped by *Nanoarchaeum equitans*. Front Microbiol 2017; 8: 1072.

19. Probst AJ, Weinmaier T, Raymann K, Perras A, Emerson JB, Rattei T, et al. Biology of a widespread uncultivated archaeon that contributes to carbon fixation in the subsurface. Nat Commun 2014; 5: 5497.

20. Giannone RJ, Huber H, Karpinets T, Heimerl T, Küper U, Rachel R, et al. Proteomic characterization of cellular and molecular processes that enable the *Nanoarchaeum equitans*- i relationship. PloS One 2011; 6: e22942.

