## Supplementary material for "An archaeal symbiont-host association from the deep terrestrial subsurface"

Content:

1. Supplementary Methods
2. Supplementary Tables
3. Supplementary Figure
4. References

### 1. Supplementary methods

#### *Phylogenetic analysis*

Hmm profiles of the 37 maker genes used by the phylosift v1.01 software [1, 2] were queried against a representative set of archaeal genomes from NCBI (182) and the GOLD database (3) as well as against the huberarchaeal population genome (for details on genomes please see Supplementary Data file 1) using the phylosift search mode [1, 2]. Protein sequences corresponding to each of the maker genes were subsequently aligned using mafft-linsi v7.407 [3] and trimmed with BMGE-1.12 (parameters: -t AA -m BLOSUM30 -h 0.55) [4]. All single protein alignments were subjected to phylogenetic analyses using IQ-tree v1.6.5 with the LG model and the tree topologies were visually inspected (iqtree -s alignment.phy -m LG -nt AUTO -bb 1000 -alrt 1000). This allowed us to select 34 proteins, in which major archaeal taxa formed monophyletic clusters suggesting that the corresponding genes were minimally affected by horizontal gene transfer. The 34 aligned and trimmed protein families were concatenated using catfasta2.phyml (https://github.com/nylander/catfasta2phyml) (alignment i) and recoded into four SR4 categories [5] (AGNPST = A, CHWY = T, DEKQR = C, FILMV = G) (alignment ii). Both alignments were subjected to maximum likelihood analyses as implemented by IQTREE (v1.6.5) under the mixture models LG+C60+F+R (alignment i, *Fig. 1A*) and the user defined C60SR4 model (alignment ii, Supplementary Data File 3) [6]. Support values were estimated using a SH-like approximate likelihood ratio test [7] and ultrafast bootstraps [8].

Furthermore, a smaller set of archaeal taxa was selected (which comprised in addition to Ca. *H. crystalense* 121 archaeal genomes from NCBI and two from the GOLD database) (Supplementary Data file 1) for Bayesian phylogenetic inferences based on the same set of 34 marker genes, which were aligned, concatenated and recoded as described above. Bayesian phylogenetic trees were inferred for the untreated and recoded alignment using the CAT-GTR model as implemented in PhyloBayes MPI v.1.7a [9]. For each alignment, a consensus tree was generated from two chains (sampling every fifth chain) once the maxdif had dropped below 0.15 (25% of the chains were discarded as burn-in).

NOTE: While the placement of Ca. *H. crystalense* within DPANN is supported in all analyses, the exact position of this genome relative to other DPANN clades needs to be further evaluated once additional genomes of members of this group are being available.

***Pangenome analysis***

For generating a comprehensive pangenome for detailed metabolic analyses, we used all available genomes from Ca. *Altiarchaeum hamiconexum* CG and Ca. *Huberiarchaeum chrystalense* [10]. This dataset encompassed at least ten individual genomes each, either from metagenomes or single-cell genomes [10]. We clustered [11] the predicted proteins of all genomes from metagenomes and single-cell genomes of either the host or the symbiont at 100% amino acid identity and selected the scaffolds of the centroids to define the pangenome. This was necessary as genomes of both the host and the symbiont were highly fragmented [10]. Some genetic redundancy in the pangenome, which stems from the fact that some scaffolds with centroids also carried non-centroid genes that had a 100% match in the dataset, did not interfere with the analysis. The data was uploaded and annotated via Genoscopes platform MAGE [12], which will be also used to make the pangenomes publicly available. Pathways were inferred using a combination of KEGG KOs [13] and final annotations from the MAGE platform. Some pathways were manually investigated and curated by inferring metabolic predictions for specific enzymes whose similarities are displayed in Supplementary Table 4 (manual annotation).

***Co-occurrence analysis***

To determine the co-occurrence analysis of Ca. *A. hamiconexum* CG and Ca. *H. crystalense* across 65 publicly available metagenome samples from Crystal Geyser [10, 15, 16], we mapped the quality filtered reads from these samples against the currently best genomes of the two organisms available from this ecosystem (quality was based on genome completeness, and N50). Reads were filtered for three mismatches (equals 98% error rate in a 150-bps read) and the coverage of each genome in each sample was determined using the iRep software [17]. Relative abundances were normalized based on the number of reads in each sample and analyzed using a Pearson correlation in the R software package [18].

***Fluoresence in situ hybridization (FISH)***

Groundwater fluids from the Crystal Geyser were filtered onto carbon coated 0.2-µm filters [especially designed for fluorescence microscopy (Millipore, Burlington, USA)] until the filters clogged (~60 ml). The samples were taken during the minor eruption phase of Crystal Geyser, during which the Altiarchaeota and Huberarchaeota abundance was the greatest [10]. Afterwards, the cells were fixed with 3% formaldehyde (16% formaldehyde (w/v) diluted in 1xPBS) for one hour at room temperature. Fixed cells were washed three times with 1xPBS. The fixed cells were stored at room temperature.

A specific probe for the 16S rRNA gene sequence of Ca. *H. crystalense* was designed using the ARB software package [19]. The probe was specific to the lineage due to the highly diverging 16S rRNA gene and did not match any other 16S rRNA gene sequences in Crystal Geyser (with less than four mismatches; HUB1206 5'-GCCCTAGACATTCGGACC-3'). The probe was 5' and 3' labeled with Cy3, while the already established probe for Altiarchaeota [20] was dual-labeled with the fluorescence-dye Atto488 (biomers.net, Ulm, Germany).

The FISH protocol was carried out as performed earlier for Altiarchaeota [20] using 0.005% SDS (w/v) and 15% deionized formamide (v/v) [21] and a probe concentration of 1 pmol/µL). Cells were counter-stained with 5 µL DAPI staining solution (DAPI solution: 50 mM EDTA pH 8.0, 0.6M Na-acetate pH 4,7, 20 ng/mL DAPI solution; 1:15 diluted in washing buffer) for 4 min at RT. The samples were examined using a Zeiss microscope (Carl Zeiss, Jena, Germany) with appropriate filter sets for Cy3, DAPI and Atto488 and a 100x Plan Apochromat objective.

The negative control for the Ca. *H. crystalense* probe was performed on biofilm samples collected from the Islinger Mühlbach, where Altiarchaeota show an extremely high abundance (~90% of the community) and of which a detailed metagenome exists [4]. The conditions for FISH were identical as described above. The probe showed no hybridization signal with Altiarchaeota or any other organisms (other Archaea or Bacteria) in the ecosystem (Figure 1-VI) displaying the high specificity of the probe (as determined *in silico*) and no cross-hybridizations with Altiarchaeota or other Bacteria in the sample.

### 2. Supplementary Tables

**Table S1** | Gene products of Ca. *A. hamiconexum* CG that could produce substrates for the metabolism of Ca. *H. chrystalense* as displayed in Figure 2. Enzymes in grey have been re-annotated manually (see Supplementary Methods and Table S3).

| No. in Fig. 2 | Label | Gene | Gene product | Product potentially used by Huberiarchaeum |
|---|---|---|---|---|
| 1 | ALTICG_v1_1430002 | pdxT | pyridoxal 5'-phosphate synthase | PLP |
| 1 | ALTICG_v1_11110003 | pdxT | pyridoxal 5'-phosphate synthase (fragment) | |
| 1 | ALTICG_v1_450003 | pdxS | pyridoxal 5'-phosphate synthase | |
| 1 | ALTICG_v1_15950001 | pdxS | pyridoxal 5'-phosphate synthase (modular protein) | |
| 2 | ALTICG_v1_120002 | ribB | 3,4-dihydroxy-2-butanone 4-phosphate synthase | FAD |
| 3 | ALTICG_v1_2080006 | | 6,7-dimethyl-8-ribityllumazine synthase (modular protein) | |
| 4 | ALTICG_v1_220004 | ribC | riboflavin synthase | |
| 4 | ALTICG_v1_15930002 | ribC | riboflavin synthase (fragment) | |
| 5 | ALTICG_v1_10003 | | putative CTP-dependent riboflavin kinase | |
| 6 | ALTICG_v1_800016 | ribL | FAD synthase | |
| 7 | ALTICG_v1_14400005 | | putative 5-amino-6-(5-phosphoribosylamino)uracil reductase | |
| 8 | ALTICG_v1_8770004 | | Diaminohydroxyphosphoribosylaminopyrimidine reductase | |
| 9 | ALTICG_v1_14400005 | | 2,5-diamino-6-ribosylamino-4(3H)-pyrimidinone 5'-phosphate reductase | |
| 10 | ALTICG_v1_780003 | nadX | putative aspartate dehydrogenase | $NAD(P)^+$ |
| 11 | ALTICG_v1_1640001 | nadA | quinolinate synthase A | |
| 12 | ALTICG_v1_2270011 | nadC | putative nicotinate-nucleotide pyrophosphorylase [carboxylating] | |
| 13 | ALTICG_v1_1540004 | | nicotinamide nucleotide adenylyltransferase | |
| 14 | ALTICG_v1_990003 | nadE | $NH_3$-dependent $NAD^+$ synthetase | |
| 15 | ALTICG_v1_7580003 | nadK | $NAD^+$ kinase | |
| 16 | ALTICG_v1_16620001 | | Nicotinate-nucleotide-dimethylbenzimidazole phosphoribosyltransferase (fragment) | |
| 17 | ALTICG_v1_10940006 | | GTP cyclohydrolase MptA | methylene-THF |
| 18 | ALTICG_v1_17060006 | | putative alkaline phosphatase | |
| 19 | ALTICG_v1_1450017 | trxB | Thioredoxin reductase | |
| 19 | ALTICG_v1_18720001 | | Thioredoxin reductase (fragment) | methylene-THF |
| 20 | ALTICG_v1_15630006 | mptE | 6-hydroxymethyl-7,8-dihydropterin pyrophosphokinase | |
| 21 | ALTICG_v1_9730008 | | Dihydropteroate synthase | |
| 22 | ALTICG_v1_14240002 | | putative dihydrofolate synthase/tetrahydrofolate synthase | |
| 23 | ALTICG_v1_2070006 | | GTP cyclohydrolase I FolE2 | |
| 24 | ALTICG_v1_15240002 | | putative anthranilate/para-aminobenzoate synthase component I | |
| 25 | ALTICG_v1_12630003 | | putative aminodeoxychorismate lyase | |
| 26 | ALTICG_v1_11410004 | | putative folylpolyglutamate synthase | |
| 27 | ALTICG_v1_2120011 | glyA | serine hydroxymethyltransferase | |
| 28 | ALTICG_v1_11270005 | | putative bifunctional NADP-dependent methylenetetrahydromethanopterin dehydrogenase /methylenetetrahydrofolate dehydrogenase | |
| 28 | ALTICG_v1_1440001 | | Methylenetetrahydrofolate dehydrogenase | |
| 29 | ALTICG_v1_1800003 | folD | bifunctional methylenetetrahydrofolate dehydrogenase/methenyltetrahydrofolate cyclohydrolase | |
| 29 | ALTICG_v1_11270006 | | putative methenyltetrahydromethanopterin cyclohydrolase | |
| 29 | ALTICG_v1_11270006 | | methenyltetrahydromethanopterin cyclohydrolase (fragment) | |
| 30 | ALTICG_v1_350005 | purU | Formyltetrahydrofolate deformylase | |
| 31 | ALTICG_v1_14850007 | metF | methylenetetrahydrofolate reductase | |
| 32 | ALTICG_v1_12010007 | | Putative tungsten-containing formylmethanofuran dehydrogenase subunit A | |

| | | | | |
|---|---|---|---|---|
| 32 | ALTICG_v1_14350006 | | putative molybdenum-containing formylmethanofuran dehydrogenase 1 subunit C | |
| 32 | ALTICG_v1_390006 | fwdB | tungsten-containing formylmethanofuran dehydrogenase 2 subunit B | |
| 32 | ALTICG_v1_10970017 | | putative tungsten-containing formylmethanofuran dehydrogenase 2 subunit C | |
| 32 | ALTICG_v1_12750017 | | formylmethanofuran dehydrogenase subunit C | |
| 32 | ALTICG_v1_12010009 | | Putative tungsten-containing formylmethanofuran dehydrogenase subunit D | |
| 33 | ALTICG_v1_790002 | ffsA | Formylmethanofuran-tetrahydromethanopterin formyltransferase | |
| 34 | ALTICG_v1_9060004 | | putative Adenylate cyclase | |
| 35 | ALTICG_v1_10330002 | | putative dihydromethanopterin reductase | |
| 36 | ALTICG_v1_17310002 | | putative acetyl-CoA synthetase | |
| 37 | ALTICG_v1_17800001 | | putative CO dehydrogenase/acetyl-CoA synthase subunit alpha 1 | |
| 37 | ALTICG_v1_19430001 | | putative CO dehydrogenase/acetyl-CoA synthase subunit alpha 2 | |
| 37 | ALTICG_v1_11000009 | | CO dehydrogenase/acetyl-CoA synthase subunit beta 2 | |
| 37 | ALTICG_v1_10980002 | | putative CO dehydrogenase/acetyl-CoA synthase subunit gamma | |
| 37 | ALTICG_v1_110003 | cdhD | CO dehydrogenase/acetyl-CoA synthase subunit delta | Acetyl-CoA for molecular building blocks |
| 38 | ALTICG_v1_16180001 | | formate dehydrogenase, nitrate-inducible, iron-sulfur subunit (fragment) | |
| 38 | ALTICG_v1_17400007 | | putative formate dehydrogenase subunit alpha | |
| 38 | ALTICG_v1_16180001 | | putative formate dehydrogenase subunit beta | |
| 38 | ALTICG_v1_1900004 | | formate dehydrogenase family accessory protein FdhD | |
| 39 | ALTICG_v1_12180007 | | gamma carbonic anhydrase family protein | |
| 39 | ALTICG_v1_12180007 | | carbonic anhydrase | |
| 40 | ALTICG_v1_12340001 | | putative lactate dehydrogenase | |
| 41 | ALTICG_v1_1540003 | porA | pyruvate ferredoxin oxidoreductase subunit alpha/ pyruvate synthase subunit porA | |
| 41 | ALTICG_v1_14030010 | porB | putative pyruvate ferredoxin oxidoreductase subunit beta/ pyruvate synthase subunit porB | |
| 41 | ALTICG_v1_14030010 | porB | putative pyruvate ferredoxin oxidoreductase subunit beta/ pyruvate synthase subunit porB (fragment) | |
| 41 | ALTICG_v1_8190001 | porC | pyruvate ferredoxin oxidoreductase subunit gamma/ pyruvate synthase subunit porC | |
| 41 | ALTICG_v1_11490003 | porD | pyruvate ferredoxin oxidoreductase subunit delta/ pyruvate synthase subunit porD | |
| 42 | ALTICG_v1_570001 | ppsA | phosphoenolpyruvate synthase | |
| 43 | ALTICG_v1_2330001 | eno | enolase | |
| 44 | ALTICG_v1_2270003 | apgM | 2,3-bisphosphoglycerate-independent phosphoglycerate mutase 1 | |
| 44 | ALTICG_v1_10720019 | | 2,3-bisphosphoglycerate-independent phosphoglycerate mutase 1 (fragment) | |
| 44 | ALTICG_v1_2490001 | | phosphoglycerate mutase | DHAP G1P |
| 45 | ALTICG_v1_5680001 | pgk | phosphoglycerate kinase | |
| 45 | ALTICG_v1_1630005 | | Phosphoglycerate kinase (fragment) | |
| 46 | ALTICG_v1_50009 | gap | glycerinaldehyde-3-phosphate dehydrogenase | |
| 47 | ALTICG_v1_450012 | tpiA | triosephosphate isomerase | |
| 48 | ALTICG_v1_2870001 | fbp | bifunctional fructose-1,6-bisphosphate aldolase/phosphatase | |
| 48 | ALTICG_v1_920002 | | Fructose-1,6-bisphosphate aldolase/phosphatase (fragment) | |
| 49 | ALTICG_v1_10090002 | pfkB | putative 6-phosphofructokinase | G1P |
| 50 | ALTICG_v1_10510014 | | Putative glucose-6-phosphate isomerase | |
| 51 | ALTICG_v1_11770006 | | putative transketolase N-terminal section | |
| 51 | ALTICG_v1_6930013 | | putative transketolase C-terminal section | |
| 52 | ALTICG_v1_1160006 | tal | putative transaldolase | |
| 53 | ALTICG_v1_1180001 | rpiA | Ribose-5-phosphate-Isomerase A | |
| 54 | ALTICG_v1_1910021 | rpe | Ribulose-phosphate 3-epimerase | |
| 55 | ALTICG_v1_11640005 | | Ribose-phosphate pyrophosphokinase | |
| 56 | ALTICG_v1_1570023 | purF | Amidophosphoribosyltransferase | |
| 57 | ALTICG_v1_70016 | purD | phosphoribosylamine-glycine ligase | GMP AMP |
| 58 | ALTICG_v1_780009 | | Phosphoribosylglycinamide formyltransferase | |
| 59 | ALTICG_v1_300002 | purL | phosphoribosylformylglycinamidine synthase subunit purL | |

| | | | | |
|---|---|---|---|---|
| 59 | ALTICG_v1_ 2610002 | | phosphoribosylformylglycinamidine synthase subunit purL (fragment) | |
| 59 | ALTICG_v1_780016 | purQ | phosphoribosylformylglycinamidine synthase subunit purQ | |
| 59 | ALTICG_v1_18190001 | | phosphoribosylformylglycinamidine synthase subunit purQ (fragment) | |
| 60 | ALTICG_v1_190011 | purM | phosphoribosylformylglycinamidine cyclo-ligase | |
| 61 | ALTICG_v1_1870002 | | putative phosphoribosylaminoimidazole carboxylase | |
| 62 | ALTICG_v1_780017 | purC | phosphoribosylaminoimidazole-succinocarboxamide synthase | |
| 63 | ALTICG_v1_10780018 | purB | Adenylosuccinate lyase | |
| 64 | ALTICG_v1_11200001 | purH | Phosphoribosylaminoimidazolecarboxamide formyltransferase/ IMP cyclohydrolase | |
| 65 | ALTICG_v1_10580005 | | putative IMP dehydrogenase | |
| 66 | ALTICG_v1_10250001 | guaAB | GMP synthase [glutamine-hydrolyzing] subunit B | |
| 67 | ALTICG_v1_11170008 | purA | Adenylosuccinate synthetase | |
| 68 | ALTICG_v1_10860008 | adkA | Adenylate kinase | UTP/dUTP |
| 69 | ALTICG_v1_1460001 | ndk | nucleoside diphosphate kinase | ATP/dATP |
| 70 | ALTICG_v1_17110001 | | putative ribonucleoside triphosphate reductase | GTP/dGTP |
| 70 | ALTICG_v1_2190001 | | Anaerobic ribonucleoside-triphosphate reductase activating protein | dCTP |
| 71 | ALTICG_v1_800002 | carB | carbamoyl-phosphate synthase large chain, C-terminal section | |
| 71 | ALTICG_v1_1570010 | carB | carbamoyl-phosphate synthase large chain, N-terminal section | |
| 71 | ALTICG_v1_141001 | carA | carbamoyl-phosphate synthase small chain | |
| 72 | ALTICG_v1_2430007 | pyrB | aspartate carbamoyltransferase | |
| 72 | ALTICG_v1_2240010 | pyrI | Aspartate carbamoyltransferase regulatory chain | |
| 73 | ALTICG_v1_3840006 | pyrC | dihydroorotase | UMP |
| 75 | ALTICG_v1_1680004 | pyrD | dihydroorotate dehydrogenase B (NAD(+)), catalytic subunit | |
| 75 | ALTICG_v1_120009 | | dihydroorotate dehydrogenase electron transfer subunit | |
| 75 | ALTICG_v1_910004 | pyrE | orotate phosphoribosyltransferase | |
| 76 | ALTICG_v1_10510002 | pyrF | orotidine-5'-phosphate decarboxylase | |
| 77 | ALTICG_v1_150001 | pyrH | uridylate kinase | dUMP/UTP |
| 78 | ALTICG_v1_990013 | tmk | putative thymidylate kinase | dTTP |
| 78 | ALTICG_v1_1100004 | tmk | dTMP kinase | |
| 79 | ALTICG_v1_860003 | thyX | Flavin-dependent thymidylate synthase | dTMP |
| 80 | ALTICG_v1_16450006 | pyrG | CTP synthase | CTP |
| 80 | ALTICG_v1_16000003 | pyrG | CTP synthase (fragment) | |
| 81 | ALTICG_v1_620001 | dcd | putative dCTP deaminase | dUTP |
| 82 | ALTICG_v1_1620001 | cmk | cytidylate kinase | CTP/dCTP |
| 83 | ALTICG_v1_13890009 | nuoI | NADH-quinone oxidoreductase subunit I | |
| 83 | ALTICG_v1_1040002 | nuoB | NADH-quinone oxidoreductase subunit B | |
| 83 | ALTICG_v1_1040003 | nuoCD | NADH-quinone oxidoreductase subunit C/D | ATP |
| 83 | ALTICG_v1_13890007 | | NADH-quinone oxidoreductase subunit D (modular protein) | |
| 83 | ALTICG_v1_1040004 | nuoH | NADH-quinone oxidoreductase subunit H | |
| 83 | ALTICG_v1_2180014 | nuoK | NADH-quinone oxidoreductase subunit K | |
| 85 | ALTICG_v1_16970009 | | ADP-ribose pyrophosphatase (fragment) | |
| 86 | ALTICG_v1_15230005 | | putative adenosine specific kinase | |
| 87 | ALTICG_v1_1760006 | | putative 3-hydroxy-3-methylglutaryl-coenzyme A synthase | |
| 88 | ALTICG_v1_11110019 | | Acetyl-CoA acetyltransferase | |
| 89 | ALTICG_v1_15560016 | hmgA | hydroxymethylglutaryl-CoA reductase | IPP |
| 90 | ALTICG_v1_1070001 | mvk | mevalonate kinase | |
| 91 | ALTICG_v1_1560003 | mvaD | diphosphomevalonate decarboxylase | |
| 92 | ALTICG_v1_14030006 | | ferredoxin | ferredoxin |
| 93 | ALTICG_v1_1740003 | fni | isopentenyl-diphosphate Delta-isomerase | |
| 94 | ALTICG_v1_3480004 | | prenyltransferase | |
| 94 | ALTICG_v1_6550010 | | prenyltransferase (fragment) | |
| 95 | ALTICG_v1_10370011 | | putative short chain isoprenyl diphosphate synthase | |
| 96 | ALTICG_v1_10240007 | | putative 2-ketoisovalerate ferredoxin oxidoreductase | |
| 97 | ALTICG_v1_15060007 | | Acylneuraminate cytidylyltransferase (fragment) | Activated sugars |
| 98 | ALTICG_v1_5710001 | egsA | NAD(P)-dependent glycerol-1-phosphate dehydrogenase | |

| | | | | |
|---|---|---|---|---|
| 99 | ALTICG_v1_12830001 | | Digeranylgeranylglyceryl phosphate synthase | |
| 100 | ALTICG_v1_2830001 | | putative digeranylgeranylglycerophospholipid reductase | |
| 101 | ALTICG_v1_1720015 | uppS | Tritrans,polycis-undecaprenyl-diphosphate synthase (geranylgeranyl-diphosphate specific) | |
| 103 | ALTICG_v1_180001 | | malate dehydrogenase | |
| 104 | ALTICG_v1_2200004 | pckG | phosphoenolpyruvate carboxykinase [GTP] | |
| 105 | ALTICG_v1_14450004 | | putative fumarate hydratase class II | |
| 105 | ALTICG_v1_16010001 | | fumarate hydratase class II (fragment) | |
| 106 | ALTICG_v1_400003 | | Aconitate hydratase | |
| 107 | ALTICG_v1_1010024 | icd | Isocitrate dehydrogenase [NADP+] | |
| 108 | ALTICG_v1_1930005 | | Succinyl-CoA synthetase subunit alpha | |
| 109 | ALTICG_v1_17410006 | | Putative CoB-CoM heterodisulfide reductase subunit B | |
| 109 | ALTICG_v1_13530010 | | Heterodisulfide reductase subunit B (fragment) | |
| 109 | ALTICG_v1_13530007 | | putative CoB-CoM heterodisulfide,ferredoxin reductase subunit A | |
| 110 | ALTICG_v1_4310002 | | putative bifunctional amino acid acetyltransferase/glutamate N-acetyltransferase | |
| 111 | ALTICG_v1_11220008 | argB | acetylglutamate kinase | |
| 112 | ALTICG_v1_2420003 | argC | N-acetyl-gamma-glutamyl-phosphate reductase | |
| 112 | ALTICG_v1_17780013 | | N-acetyl-gamma-glutamyl-phosphate reductase (fragment) | |
| 113 | ALTICG_v1_2010003 | argD | acetylornithine aminotransferase | Arg |
| 114 | ALTICG_v1_14340003 | argJ | glutamate N-acetyltransferase | |
| 115 | ALTICG_v1_1470017 | argF | ornithine carbamoyltransferase | |
| 116 | ALTICG_v1_2490003 | argG | argininosuccinate synthase | |
| 116 | ALTICG_v1_15110003 | | argininosuccinate synthase (fragment) | |
| 117 | ALTICG_v1_990011 | argH | argininosuccinate lyase | |
| 117 | ALTICG_v1_14200008 | | argininosuccinate lyase (fragment) | |
| 118 | ALTICG_v1_10970009 | | Arginase | |
| 119 | ALTICG_v1_3900006 | glnA | glutamine synthetase | |
| 119 | ALTICG_v1_15030004 | | glutamine synthetase (fragment) | |
| 120 | ALTICG_v1_15790002 | phzB | Glutamine amidotransferase | |
| 121 | ALTICG_v1_2720003 | gdhA | glutamate dehydrogenase, NADP-specific | Gln |
| 121 | ALTICG_v1_16090004 | | glutamate dehydrogenase, NADP-specific (fragment) | Glu |
| 122 | ALTICG_v1_310003 | aspC | aspartate aminotransferase | Asp |
| 123 | ALTICG_v1_2560007 | ansB | asparagine synthase (Glutamine-hydrolyzing) | Asn |
| 123 | ALTICG_v1_14390004 | | asparagine synthase (fragment) | |
| 124 | ALTICG_v1_990007 | ansB | Aspartate ammonia-lyase | |
| 124 | ALTICG_v1_14450004 | | Aspartate ammonia-lyase (fragment) | Asp |
| 125 | ALTICG_v1_10220001 | | Aspartate kinase (fragment) | homoserine |
| 126 | ALTICG_v1_220003 | asd | aspartate-semialdehyde dehydrogenase | Thr |
| 127 | ALTICG_v1_1950007 | hom | Homoserine dehydrogenase | |
| 128 | ALTICG_v1_350002 | thrC | threonine synthase | |
| 129 | ALTICG_v1_11920007 | cimA | (R)-citramalate synthase | |
| 130 | ALTICG_v1_1880008 | leuC | 3-isopropylmalate dehydratase large subunit | |
| 130 | ALTICG_v1_13380003 | leuD | 3-isopropylmalate dehydratase small subunit 2 | |
| 130 | ALTICG_v1_4780014 | leuD | Isopropylmalate/citramalate isomerase small subunit | |
| 131 | ALTICG_v1_7880008 | leuB | 3-isopropylmalate dehydrogenase | |
| 132 | ALTICG_v1_15550018 | ilvB | putative acetolactate synthase large subunit | Val |
| 132 | ALTICG_v1_1010008 | ilvH | putative acetolactate synthase small subunit | Leu |
| 133 | ALTICG_v1_1910009 | ilvC | Ketol-acid reductoisomerase (NADP(+)) | Ile |
| 133 | ALTICG_v1_15580001 | | Ketol-acid reductoisomerase (NADP(+)) (fragment) | |
| 134 | ALTICG_v1_1450020 | ilvD | Dihydroxy-acid dehydratase | |
| 135 | ALTICG_v1_1720003 | ilvE | putative branched-chain-amino-acid aminotransferase | |
| 136 | ALTICG_v1_2050003 | leuA | putative 2-isopropylmalate synthase | |
| 136 | ALTICG_v1_8980008 | | 2-isopropylmalate synthase (fragment) | |
| 137 | ALTICG_v1_800012 | dapA | 4-hydroxy-tetrahydrodipicolinate synthase | Lys |

| | | | | |
|---|---|---|---|---|
| 138 | ALTICG_v1_2490005 | dapB | 4-hydroxy-tetrahydrodipicolinate reductase | |
| 139 | ALTICG_v1_760001 | dapL | LL-diaminopimelate aminotransferase | |
| 140 | ALTICG_v1_10940005 | dapF | Diaminopimelate epimerase | |
| 141 | ALTICG_v1_10050010 | lysA | Diaminopimelate decarboxylase | |
| 142 | ALTICG_v1_1080003 | pdaD | putative pyruvoyl-dependent arginine decarboxylase | |
| 143 | ALTICG_v1_10970009 | speB | agmatinase | |
| 143 | ALTICG_v1_17630001 | | Agmatinase (fragment) | |
| 144 | ALTICG_v1_120010 | hisG | ATP phosphoribosyltransferase | |
| 145 | ALTICG_v1_11110023 | | Putative phosphoribosyl-ATP pyrophosphohydrolase/ phosphoribosyl-AMP cyclohydrolase | |
| 146 | ALTICG_v1_1900017 | | Phosphoribosyl-AMP cyclohydrolase | |
| 147 | ALTICG_v1_13830005 | hisA | 1-(5-phosphoribosyl)-5-[(5-phosphoribosylamino) methylideneamino] imidazole-4-carboxamide isomerase | His |
| 148 | ALTICG_v1_10220016 | | putative imidazoleglycerol-phosphate synthase cyclase subunit | |
| 148 | ALTICG_v1_18360002 | | putative imidazoleglycerol-phosphate synthase | |
| 149 | ALTICG_v1_10220014 | | Imidazoleglycerol-phosphate dehydratase | |
| 150 | ALTICG_v1_10460002 | hisC | Histidinol-phosphate aminotransferase | |
| 151 | ALTICG_v1_1630001 | hisD | histidinol dehydrogenase | |
| 152 | ALTICG_v1_15820005 | aroB | 3-dehydroquinate synthase | |
| 153 | ALTICG_v1_10720016 | | putative 3-dehydroquinate dehydratase | |
| 154 | ALTICG_v1_1610003 | aroE | Shikimate dehydrogenase (NADP(+)) | |
| 155 | ALTICG_v1_14980003 | aroK | Shikimate kinase | |
| 156 | ALTICG_v1_12030009 | aroA | 3-phosphoshikimate 1-carboxyvinyltransferase | |
| 157 | ALTICG_v1_390001 | aroC | chorismate synthase | |
| 158 | ALTICG_v1_17100001 | | chorismate mutase (fragment) | |
| 159 | ALTICG_v1_800018 | pheA | Prephenate dehydratase | |
| 160 | ALTICG_v1_2500003 | | Prephenate dehydrogenase | Trp<br>Phe<br>Tyr |
| 160 | ALTICG_v1_12940002 | | putative arogenate/prephenate dehydrogenase (fragment) | |
| 161 | ALTICG_v1_14840010 | | aromatic amino acid aminotransferase | |
| 162 | ALTICG_v1_15810007 | mfnA | L-tyrosine/L-aspartate decarboxylase | |
| 162 | ALTICG_v1_15270005 | | L-aspartate decarboxylase (fragment) | |
| 163 | ALTICG_v1_11290002 | | Phenazine-specific anthranilate synthase component I | |
| 164 | ALTICG_v1_720002 | trpD | Anthranilate phosphoribosyltransferase | |
| 165 | ALTICG_v1_13630003 | trpC | Indole-3-glycerol phosphate synthase | |
| 166 | ALTICG_v1_2400007 | trpA | tryptophan synthase alpha chain | |
| 166 | ALTICG_v1_2260002 | trpB | tryptophan synthase beta subunit | |
| 167 | ALTICG_v1_14850006 | metE | 5-methyltetrahydropteroyltriglutamate-homocysteine methyltransferase | |
| 168 | ALTICG_v1_1740011 | mat | S-adenosylmethionine synthase | SAM |
| 168 | ALTICG_v1_17700001 | | putative Methionine adenosyltransferase | |
| 170 | ALTICG_v1_380003 | ahcY | adenosylhomocysteinase | |
| 171 | ALTICG_v1_9220013 | mqnC | Cyclic dehypoxanthine futalosine synthase | SAM |
| 172 | ALTICG_v1_2920007 | pcm | Protein-L-isoaspartate O-methyltransferase | |
| 173 | ALTICG_v1_14410003 | dphB | Diphthine synthase | |
| 174 | ALTICG_v1_16240001 | | Phosphomethylpyrimidine synthase (fragment) | SAM |
| 175 | ALTICG_v1_13910009 | | putative homoserine O-acetyltransferase | |
| 176 | ALTICG_v1_2350004 | serB | Phosphoserine phosphatase | Ser<br>Ala<br>Cys |
| 177 | ALTICG_v1_2470003 | iscS | Cysteine desulfurase | |
| 178 | ALTICG_v1_19270004 | | putative serine acetyltransferase | |
| 179 | ALTICG_v1_550003 | cobT | Nicotinate-nucleotide-dimethylbenzimidazole phosphoribosyltransferase | |
| 180 | ALTICG_v1_11320004 | | cobalt chelatase | |
| 181 | ALTICG_v1_140011 | cobD | Cobalamin biosynthesis protein CobD | |
| 182 | ALTICG_v1_140005 | | Cob(I)yrinic acid a,c-diamide adenosyltransferase | AdoCbl |
| 183 | ALTICG_v1_2500001 | | Bifunctional adenosylcobinamide kinase/adenosylcobinamide-phosphate guanylyltransferase | |
| 184 | ALTICG_v1_1810001 | cobS | Adenosylcobinamide-GDP ribazoletransferase | |

| | | | | |
|---|---|---|---|---|
| 185 | ALTICG_v1_2780010 | | putative UTP-glucose-1-phosphate uridylyltransferase | |
| 185 | ALTICG_v1_1610011 | | UTP-glucose-1-phosphate uridylyltransferase (fragment) | |
| 186 | ALTICG_v1_890014 | | putative UDP-glucose 4-epimerase | |
| 187 | ALTICG_v1_40007 | tuaD | UDP-glucose-6-dehydrogenase | |
| 188 | ALTICG_v1_40008 | | putative UDP-D-glucuronate decarboxylase | |
| 189 | ALTICG_v1_14630003 | | putative UDP-4-amino-4-deoxy-L-arabinose-oxoglutarate aminotransferase | |
| 190 | ALTICG_v1_800023 | | bifunctional phosphoglucomutase/phosphomannomutase | |
| 191 | ALTICG_v1_18750004 | xanB | mannose-6-phosphate isomerase/mannose-1-phosphate guanylyltransferase | |
| 192 | ALTICG_v1_1610007 | strD | glucose-1-phosphate thymidylyltransferase | |
| 192 | ALTICG_v1_7310001 | | glucose-1-phosphate thymidylyltransferase (fragment) | |
| 193 | ALTICG_v1_10970018 | | putative glucose-1-phosphate adenylyltransferase | |
| 194 | ALTICG_v1_1600004 | rfbF | glucose-1-phosphate cytidylyltransferase | |
| 195 | ALTICG_v1_1360008 | tpsp | Alpha,alpha-trehalose-phosphate synthase [UDP-forming] / trehalose-6-phosphate phosphatase | |
| 196 | ALTICG_v1_6330013 | gmd | GDP-D-mannose 4,6-dehydratase, NAD(P)-binding | |
| 197 | ALTICG_v1_8660002 | | GDP-L-fucose synthase (fragment) | |
| 198 | ALTICG_v1_2740002 | rmlC | dTDP-4-dehydrorhamnose 3,5-epimerase | Activated sugars G1P |
| 199 | ALTICG_v1_17460001 | | glucosamine-fructose-6-phosphate aminotransferase (fragment) | |
| 200 | ALTICG_v1_12370014 | glmM | phosphoglucosamine mutase | |
| 201 | ALTICG_v1_2100006 | glmU | UDP-N-acetylglucosamine pyrophosphorylase / Glucosamine-1-phosphate N-acetyltransferase | |
| 202 | ALTICG_v1_13920009 | | UDP-N-acetylglucosamine pyrophosphorylase / Glucosamine-1-phosphate N-acetyltransferase (fragment) | |
| 203 | ALTICG_v1_510005 | wecB | UDP-N-acetylglucosamine 2-epimerase | |
| 203 | ALTICG_v1_15990003 | | UDP-N-acetylglucosamine 2-epimerase (fragment) | |
| 204 | ALTICG_v1_14630006 | | UDP-N-acetylglucosamine 3-dehydrogenase (fragment) | |
| 205 | ALTICG_v1_18870008 | | dTDP-glucose 4,6-dehydratase (fragment) | |
| 205 | ALTICG_v1_1610005 | rmlB | dTDP-glucose 4,6 dehydratase, NAD(P)-binding | |
| 206 | ALTICG_v1_510003 | wecC | UDP-N-acetyl-D-mannosamine dehydrogenase | |
| 206 | ALTICG_v1_16820004 | | UDP-N-acetyl-D-glucosamine 6-dehydrogenase (fragment) | |
| 207 | ALTICG_v1_10760013 | | putative dTDP-4-dehydrorhamnose reductase | |
| 208 | ALTICG_v1_11690017 | pseH | UDP-4-amino-4, 6-dideoxy-N-acetyl-beta-L-altrosamine N-acetyltransferase | |
| 209 | ALTICG_v1_19330001 | amC | Undecaprenyl-phosphate 4-deoxy-4-formamido-L-arabinose transferase | |
| 210 | ALTICG_v1_1790001 | | putative endoglucanase/ cellulase/ putative peptidase M42 | |
| 211 | ALTICG_v1_60004 | | putative alpha-amylase | |
| 212 | ALTICG_v1_3110006 | | alpha-1,4 glucan phosphorylase | |
| 213 | ALTICG_v1_11570001 | | Glycogen debranching protein | |
| 214 | ALTICG_v1_11290001 | | putative glycogen [starch] synthase | glycogen |
| 215 | ALTICG_v1_3550008 | | putative CDP-4-dehydro-6-deoxy-D-gulose 4-reductase | Activated sugars |
| 216 | ALTICG_v1_10210015 | | putative Ferredoxin-NADP reductase | red. ferredoxin |
| 217 | ALTICG_v1_13890016 | arsC | Arsenate reductase | |
| 218 | ALTICG_v1_13890012 | arsM | Arsenite methyltransferase | |
| 219 | ALTICG_v1_13890017 | arsB | Arsenite resistance protein ArsB | |
| 220 | ALTICG_v1_11110015 | | putative archaeal A1AO-type ATP synthase, subunit K | |
| 220 | ALTICG_v1_14070002 | | putative archaeal A1AO-type ATP synthase, subunit B | |
| 220 | ALTICG_v1_15150001 | | putative archaeal A1AO-type ATP synthase, subunit A | |
| 220 | ALTICG_v1_10430022 | | putative archaeal A1AO-type ATP synthase, subunit D | ATP |
| 220 | ALTICG_v1_12040009 | | putative archaeal A1AO-type ATP synthase, subunit C | |
| 220 | ALTICG_v1_16550006 | | putative archaeal A1AO-type ATP synthase, subunit I | |

122

**Table S2** | Gene products of Ca. *H. chrystalense* that could potentially use substrates provided by Ca. *A. hamiconexum* as displayed in Figure 2. Enzymes in grey have been re-annotated manually (see Supplementary Methods and Table S3).

| No. In Fig. 2 | Label | Gene | Gene product | Component potentially derived from Altiarchaeum |
|---|---|---|---|---|
| 19 | HUBERARCH_v1_220006 | trxB | Thioredoxin reductase | NAD(P)H |
| 19 | HUBERARCH_v1_4870001 | | Thioredoxin reductase (fragment) | NAD(P)H |
| 69 | HUBERARCH_v1_4750008 | ndk | Nucleoside diphosphate kinase | ATP/GTP |
| 70 | HUBERARCH_v1_100006 | nrdD | anaerobic ribonucleoside triphosphate reductase | AdoCbl |
| 78 | HUBERARCH_v1_230004 | tmk | putative thymidylate kinase | ATP/GTP |
| 79 | HUBERARCH_v1_1440004 | | Putative thymidylate synthase | methylene-THF |
| 80 | HUBERARCH_v1_260014 | pyrG | CTP synthase | ATP/GTP |
| 84 | HUBERARCH_v1_110006 | | putative CMP/dCMP deaminase zinc-binding protein, dCMP deaminase | $Zn^{2+}$ |
| 93 | HUBERARCH_v1_370010 | idi | Isopentenyl-diphosphate delta-isomerase 2 | |
| 94 | HUBERARCH_v1_80001 | | Geranylgeranyl diphosphate synthase | |
| 98 | HUBERARCH_v1_3540004 | egsA | NAD(P)-dependent glycerol-1-phosphate dehydrogenase | NAD(P)H |
| 98 | HUBERARCH_v1_1470004 | | NAD(P)-dependent glycerol-1-phosphate dehydrogenase (fragment) | NAD(P)H |
| 100 | HUBERARCH_v1_1430005 | | putative digeranylgeranylglycerophospholipid reductase | reduced ferredoxin |
| 102 | HUBERARCH_v1_800015 | ytsJ | putative NAD-dependent malic enzyme 4 | NAD(P)H |
| 103 | HUBERARCH_v1_10014 | | putative malic protein NAD-binding protein, malate dehydrogenase (Oxaloacetate-decarboxylating) | NAD(P)H |
| 103 | HUBERARCH_v1_10014 | ytsJ | putative malate dehydrogenase | NAD(P)H |
| 118 | HUBERARCH_v1_2100015 | | arginase | |
| 122 | HUBERARCH_v1_100004 | | putative aspartate aminotransferase | PLP |
| 123 | HUBERARCH_v1_110022 | | putative asparagine synthase | |
| 143 | HUBERARCH_v1_3660016 | | putative agmatinase | |
| 159 | HUBERARCH_v1_100003 | | Prephenate dehydratase | |
| 166 | HUBERARCH_v1_1170012 | | putative tryptophan synthase alpha chain | PLP |
| 169 | HUBERARCH_v1_4370002 | | DNA (Cytosine-5-)-methyltransferase | SAM |
| 172 | HUBERARCH_v1_950023 | | Protein-L-isoaspartate O-methyltransferase | SAM |
| 173 | HUBERARCH_v1_220013 | dph | Diphthine synthase | SAM |
| 177 | HUBERARCH_v1_630002 | iscS | cysteine desulfurylase | PLP |
| 186 | HUBERARCH_v1_860003 | | putative UDP-glucose 4-epimerase | |
| 192 | HUBERARCH_v1_2660001 | strD | glucose-1-phosphate thymidylyltransferase | dTTP |
| 193 | HUBERARCH_v1_3470001 | | putative glucose-1-phosphate adenylyltransferase | ATP |
| 194 | HUBERARCH_v1_520002 | rfbF | Glucose-1-phosphate cytidylyltransferase | CTP |
| 196 | HUBERARCH_v1_520006 | | GDP-mannose 4,6-dehydratase | |
| 198 | HUBERARCH_v1_520004 | rfbC | dTDP-4-dehydrorhamnose 3,5-epimerase | |
| 198 | HUBERARCH_v1_860002 | rmlC | dTDP-4-dehydrorhamnose 3,5-epimerase | |
| 203 | HUBERARCH_v1_360004 | | UDP-N-acetylglucosamine 2-epimerase (Non-hydrolyzing) | |
| 203 | HUBERARCH_v1_2160021 | | UDP-N-acetylglucosamine 2-epimerase (fragment) | |
| 205 | HUBERARCH_v1_960005 | rmlB | dTDP-glucose 4,6-dehydratase, NAD(P)-binding | |
| 207 | HUBERARCH_v1_2660002 | | putative dTDP-4-dehydrorhamnose reductase | |
| 210 | HUBERARCH_v1_1780008 | | putative endoglucanase/ cellulose/ putative peptidase M42 | |
| 215 | HUBERARCH_v1_2860003 | | CDP-4-dehydro-6-deoxy-D-gulose 4-reductase | |
| 221 | HUBERARCH_v1_4290002 | | DNA alkylation repair protein | |
| | HUBERARCH_v1_4600012 | | DNA helicase UvrD | |
| | HUBERARCH_v1_4520009 | radA | DNA repair and recombination protein RadA | |
| | HUBERARCH_v1_4470025 | radB | DNA repair and recombination protein RadB | |
| | HUBERARCH_v1_4350016 | lig | DNA ligase | |

| | | | |
|---|---|---|---|
| 222 | HUBERARCH_v1_4570037 | lon | Archaeal Lon protease |
| | HUBERARCH_v1_2850003 | htpX | Protease HtpX homolog |
| | HUBERARCH_v1_3710005 | | Proteasome endopeptidase complex |
| | HUBERARCH_v1_4860003 | pan | Proteasome-activating nucleotidase |
| | HUBERARCH_v1_4470021 | rrp | Exosome complex component Rrp41 |
| | HUBERARCH_v1_4350010 | | Metallopeptidase |
| 223 | HUBERARCH_v1_4280019 | cdc6 | ORC1-type DNA replication protein 1 |
| | HUBERARCH_v1_4360019 | | Replication factor C large subunit (fragment) |
| | HUBERARCH_v1_4610004 | rfcS | Replication factor C small subunit |
| | HUBERARCH_v1_240025 | polC | DNA polymerase II large subunit |
| | HUBERARCH_v1_1400015 | polB | putative DNA polymerase II small subunit |
| 224 | HUBERARCH_v1_4780003 | rpoE | DNA-directed RNA polymerase subunit E' |
| | HUBERARCH_v1_150003 | rpoH | DNA-directed RNA polymerase subunit H |
| | HUBERARCH_v1_4640001 | | DNA-directed RNA polymerase subunit N |
| | HUBERARCH_v1_4320008 | | DNA-directed RNA polymerase subunit P (fragment) |
| | HUBERARCH_v1_4380029 | spt | Transcription elongation factor Spt4 |
| | HUBERARCH_v1_4470024 | | Transcription elongation factor Spt5 |
| | HUBERARCH_v1_520007 | | NarL family transcriptional regulator |
| | HUBERARCH_v1_4280033 | tfs | Transcription factor S |
| | HUBERARCH_v1_850001 | | Transcriptional regulator |
| 225 | HUBERARCH_v1_4420009 | | Protein translation factor SUI1 homolog |
| | HUBERARCH_v1_4710005 | | Ribosome assembly factor SBDS |
| | HUBERARCH_v1_4820004 | | Translation initiation factor IF-1A |
| | HUBERARCH_v1_1800003 | | Translation initiation factor IF-2 subunit beta |
| | HUBERARCH_v1_1180003 | | putative translation initiation factor 2 subunit gamma |
| | HUBERARCH_v1_4430014 | eif5A | Translation initiation factor IF-5A |
| | HUBERARCH_v1_4390027 | prf | Peptide chain release factor 1 |
| | HUBERARCH_v1_4390022 | tuf | Elongation factor 1-alpha |
| | HUBERARCH_v1_4520006 | fusA | Elongation factor 2 |
| | HUBERARCH_v1_4690004 | | Nascent polypeptide-associated complex protein |
| | HUBERARCH_v1_3060007 | iscU | scaffold protein |
| | HUBERARCH_v1_4500005 | alaS | Alanine--tRNA ligase |
| | HUBERARCH_v1_3590030 | | Alanyl-tRNA editing protein AlaX |
| | HUBERARCH_v1_3590030 | | Alanyl-tRNA editing protein AlaX-M (fragment) |
| | HUBERARCH_v1_4420021 | argS | Arginine--tRNA ligase |
| | HUBERARCH_v1_70012 | aspS | Aspartate--tRNA(Asp) ligase |
| | HUBERARCH_v1_4640004 | gltX | Glutamate--tRNA ligase |
| | HUBERARCH_v1_4390026 | glyQS | Glycine--tRNA ligase |
| | HUBERARCH_v1_4720003 | hisS | Histidine--tRNA ligase 1//Histidine--tRNA ligase |
| | HUBERARCH_v1_4730001 | | Isoleucine--tRNA ligase |
| | HUBERARCH_v1_4570039 | leuS | Leucine--tRNA ligase |
| | HUBERARCH_v1_3310003 | lysS | Lysine--tRNA ligase |
| | HUBERARCH_v1_4060001 | metG | Methionine--tRNA ligase |
| | HUBERARCH_v1_4550008 | | O-phospho-L-seryl-tRNA:Cys-tRNA synthase 1 (fragment)//O-phospho-L-seryl-tRNA:Cys-tRNA synthase |
| | HUBERARCH_v1_4550005 | | O-phosphoserine--tRNA ligase |
| | HUBERARCH_v1_4470010 | | Phenylalanine--tRNA ligase subunit beta |
| | HUBERARCH_v1_4740002 | proS | Proline--tRNA ligase |
| | HUBERARCH_v1_4630008 | serS | Serine--tRNA ligase |
| | HUBERARCH_v1_4380006 | thrS | Threonine--tRNA ligase |
| | HUBERARCH_v1_4650006 | trpS | Tryptophan--tRNA ligase |
| | HUBERARCH_v1_4420012 | tyrS | Tyrosine--tRNA ligase |
| | HUBERARCH_v1_3850006 | valS | Valine--tRNA ligase |
| | HUBERARCH_v1_4470019 | cca | CCA tRNA nucleotidyltransferase |

| | | | |
|---|---|---|---|
| | HUBERARCH_v1_4670003 | taw | S-adenosyl-L-methionine-dependent tRNA 4-demethylwyosine synthase//tRNA 4-demethylwyosine synthase (AdoMet-dependent) |
| | HUBERARCH_v1_4390023 | kae | tRNA N6-adenosine threonylcarbamoyltransferase//N(6)-L-threonylcarbamoyladenine synthase |
| | HUBERARCH_v1_1990005 | taw | tRNA(Phe) 7-((3-amino-3-carboxypropyl)-4-demethylwyosine(37)-N(4))-methyltransferase |
| | HUBERARCH_v1_4390009 | | tRNA (cytidine(56)-2'-O)-methyltransferase |
| | HUBERARCH_v1_4550008 | | O-phospho-L-seryl-tRNA:Cys-tRNA synthase 1 (fragment)//O-phospho-L-seryl-tRNA:Cys-tRNA synthase |
| | HUBERARCH_v1_4280013 | map | Methionine aminopeptidase |
| 226 | HUBERARCH_v1_4360016 | | Cell division cycle protein 48 homolog AF_1297 |
| | HUBERARCH_v1_4400024 | | Cell division protein DedD |
| | HUBERARCH_v1_4670006 | ftsZ | Cell division protein FtsZ 1 |
| | HUBERARCH_v1_4390010 | ftsZ | Cell division protein FtsZ 2 |
| | HUBERARCH_v1_4350004 | | Chromosomal protein MC1c |
| | HUBERARCH_v1_4390019 | topA | DNA topoisomerase 1 |
| | HUBERARCH_v1_1400025 | | Histone |
| | HUBERARCH_v1_2580031 | | Histone (fragment) |
| | HUBERARCH_v1_3900003 | | putative transposase |
| 227 | HUBERARCH_v1_3390001 | | DNA adenine methylase |
| | HUBERARCH_v1_4510001 | | Site-specific DNA-methyltransferase |
| | HUBERARCH_v1_3390002 | | RNA methyltransferase |
| 228 | HUBERARCH_v1_4310007 | | Beta-CASP ribonuclease aCPSF1 |
| | HUBERARCH_v1_1160001 | | Putative RNA-binding protein RbpB (fragment) |
| | HUBERARCH_v1_4670005 | | putative snRNP Sm-like protein |
| | HUBERARCH_v1_4510002 | | Restriction endonuclease subunit R |
| | HUBERARCH_v1_3220009 | | Ribonuclease |
| | HUBERARCH_v1_4380031 | rnhB | Ribonuclease H |
| | HUBERARCH_v1_3750016 | rnp | Ribonuclease P protein component 2//Ribonuclease P |
| | HUBERARCH_v1_4430009 | rnz | Ribonuclease Z |
| | HUBERARCH_v1_4660001 | | RNA 2',3'-cyclic phosphodiesterase |
| | HUBERARCH_v1_4320005 | flpA | Fibrillarin-like rRNA/tRNA 2'-O-methyltransferase |
| | HUBERARCH_v1_1100002 | | Radical SAM protein |
| | HUBERARCH_v1_3710008 | | Restriction endonuclease |
| | HUBERARCH_v1_300001 | | Restriction endonuclease subunit M |
| | HUBERARCH_v1_4470022 | | RNA-binding protein |
| | HUBERARCH_v1_2860001 | | SAM-dependent methyltransferase |
| | HUBERARCH_v1_3390007 | | Type I restriction endonuclease subunit M |
| | HUBERARCH_v1_2730001 | | Type I restriction endonuclease subunit M |
| | HUBERARCH_v1_4370001 | | Type II restriction endonuclease |
| 229 | HUBERARCH_v1_3210001 | | Glycosyl transferase family |
| | HUBERARCH_v1_4770003 | | Glycosyl transferase family 1 |
| | HUBERARCH_v1_3320001 | | Glycosyl transferase family 2 |
| | HUBERARCH_v1_4780002 | | Glycosyl transferase family 4 |
| | HUBERARCH_v1_3650003 | agl | Low-salt glycan biosynthesis nucleotidyltransferase Agl11 |

**Table S3** | Cutoffs and taxonomy of the best hits of manually annotated enzymes (in grey in Table S1 and Table S2).

| Number in Table S2 | % identity | taxonomy of the respective hit |
|---:|---:|---|
| 5 | 77.95 | groundwater genome |
| 7 | 47.70 | Methanobacterium bryantii |
| 18 | 82.23 | HGW-Altiarchaeales-1 |
| 22 | 86.84 | groundwater metagenome |
| 24 | 40.46 | Burkholderia lata (strain ATCC 17760 / DSM 23089 / LMG 22485 / NCIMB 9086 / R18194 / 383) |
| 25 | 44.33 | Bacillus subtilis (strain 168) |
| 26 | 79.49 | groundwater metagenome |
| 28 | 39.66 | Methylobacterium extorquens (strain ATCC 14718 / DSM 1338 / JCM 2805 / NCIMB 9133 / AM1) |
| 29 | 90.57 | groundwater metagenome |
| 32 | 46.55-87.02 | groundwater metagenome/ *Methanothermobacter wolfeii* |
| 35 | 38.46 | *Methanocaldococcus jannaschii* (strain ATCC 43067 / DSM 2661 / JAL-1 / JCM 10045 / NBRC 100440) |
| 36 | 68.75 | Candidatus Entotheonella factor |
| 37 | 66.35-89.89 | groundwater metagenome |
| 38 | 90.00-91.08 | groundwater metagenome |
| 41 | 56.34 | Methanococcus maripaludis X1 |
| 50 | 83.04 | groundwater metagenome |
| 61 | 91.89 | groundwater metagenome |
| 70 | 91.60 | Candidatus Altiarchaeales archaeon A3 |
| 86 | 70.27 | *Methanomassiliicoccales* archaeon PtaU1.Bin124 |
| 87 | 90.31 | groundwater metagenome |
| 95 | 86.24 | groundwater metagenome |
| 96 | 54.67 | *Thermococcus guaymasensis* |
| 100 | 80.08 | groundwater metagenome |
| 105 | 55.12 | *Thermoflexus hugenholtzii* JAD2 |
| 109 | 43.93-87.5/ | groundwater metagenome/ *Methanococcus maripaludis* (strain S2 / LL) |
| 145 | 50.00 | *Escherichia coli* |
| 148 | 52.59-50.00 | *Rhizobium etli/Thermoanaerobacter ethanolicus* |
| 153 | 94.23 | HGW-Altiarchaeales-2 |
| 175 | 45.69 | *Methanococcoides burtonii* (strain DSM 6242 / NBRC 107633 / OCM 468 / ACE-M) |
| 178 | 78.08 | *Aciduliprofundum sp.* (strain MAR08-339) |
| 188 | 49.64 | *Nicotiana tabacum* |
| 189 | 42.03 | *Salmonella typhimurium* (strain LT2 / SGSC1412 / ATCC 700720) |
| 193 | 84.44 | groundwater metagenome |
| 207 | 85.21 | groundwater metagenome |
| 210 | 47.48 | Methanococcus maripaludis |
| 214 | 85.95 | groundwater metagenome |
| 215 | 89.80 | archaeon (Candidatus Huberarchaea) CG03_land_8_20_14_0_80_31_114 |
| 216 | 86.17 | groundwater metagenome |
| 220 | 80.68-93.84 | groundwater genome |

| Number in Table S3 | % identity | taxonomy of the respective hit |
|---:|---:|---|
| 79 | 38.27 | Parcubacteria group bacterium GW2011_GWF2_43_11 |
| 84 | 61.25 | Berkelbacteria bacterium GW2011_GWE1_39_12 |
| 100 | 66.27 | groundwater metagenome |
| 103 | 55.36 | Candidatus Peregrinibacteria bacterium GW2011_GWF2_43_17 |
| 143 | 53.70 | *Methanobacterium* sp. PtaU1.Bin242 |
| 166 | 43.42 | *Minicystis rosea* |
| 186 | 45.71 | *Arabidopsis thaliana* |
| 193 | 90.12 | groundwater metagenome |

| | | |
|---|---|---|
| 207 | 78.69 | groundwater metagenome |
| 210 | 48.99 | *Methanobrevibacter olleyae* |
| 225 | 53.33 | Candidatus Pacearchaeota archaeon |
| 226 | 73.10 | Candidatus Methanoperedens nitroreducens |

**Fig. S1** | Ratios of relative abundances of Altiarchaeota and Huberarchaeota (based on stringent mapping of reads) across all samples. With the exception of a few smaller filters (0.1 µm and 0.2 µm), the ratio is about 11/1, i.e. eleven Ca. *Altiarchaeota* per one Ca. *Huberiarchaeum*.
For some filters with a 0.1 µm and 0.2 µm pore size, the number of Huberarchaeota per Altiarchaeota increase, indicating that *Ca.* Huberiarchaeum has a small cell size [10] and thus passes through the larger filters while *Ca.* Altiarchaeum does not.

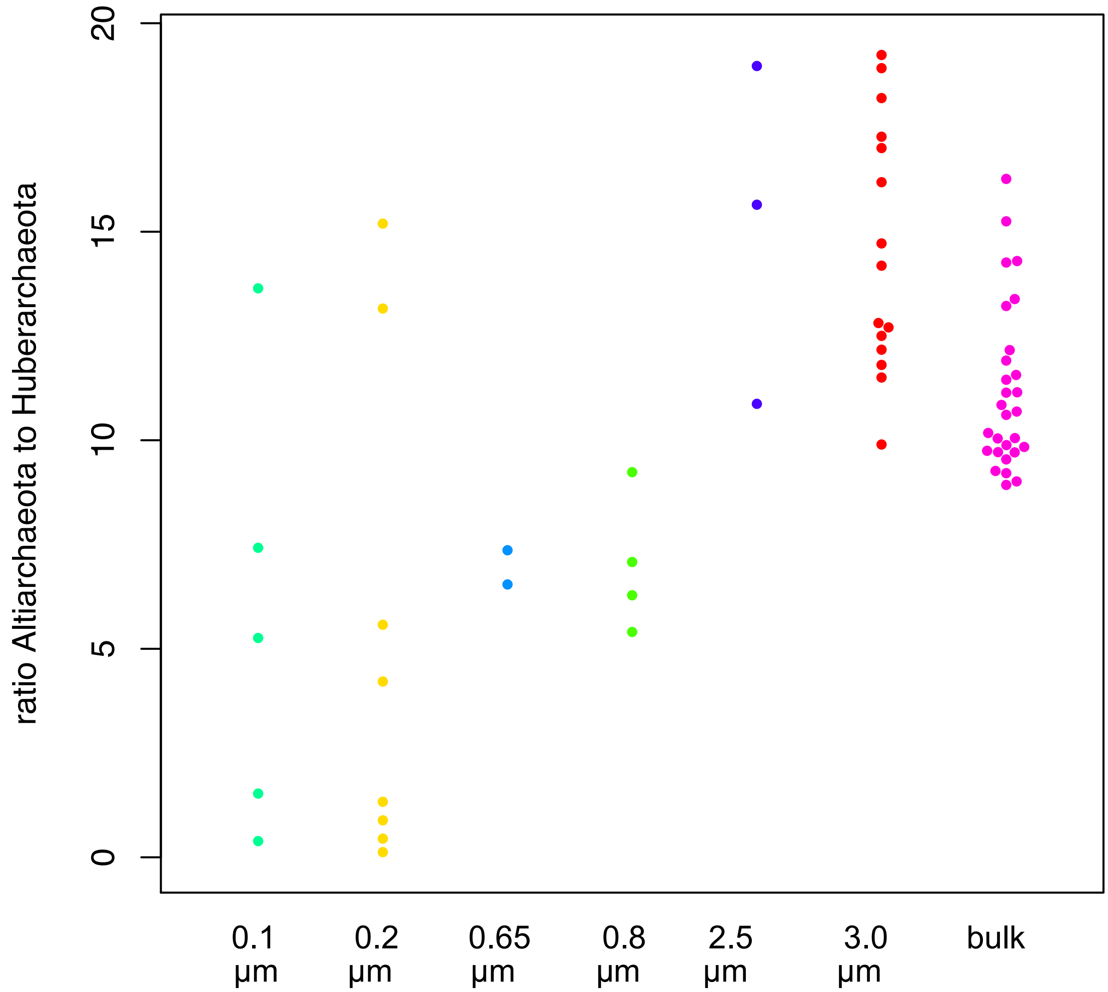

**Fig. S2** | Detailed metabolic reactions of predicted enzymes from Supplementary Table 1 and 2. Summary is provided in main Figure 2.

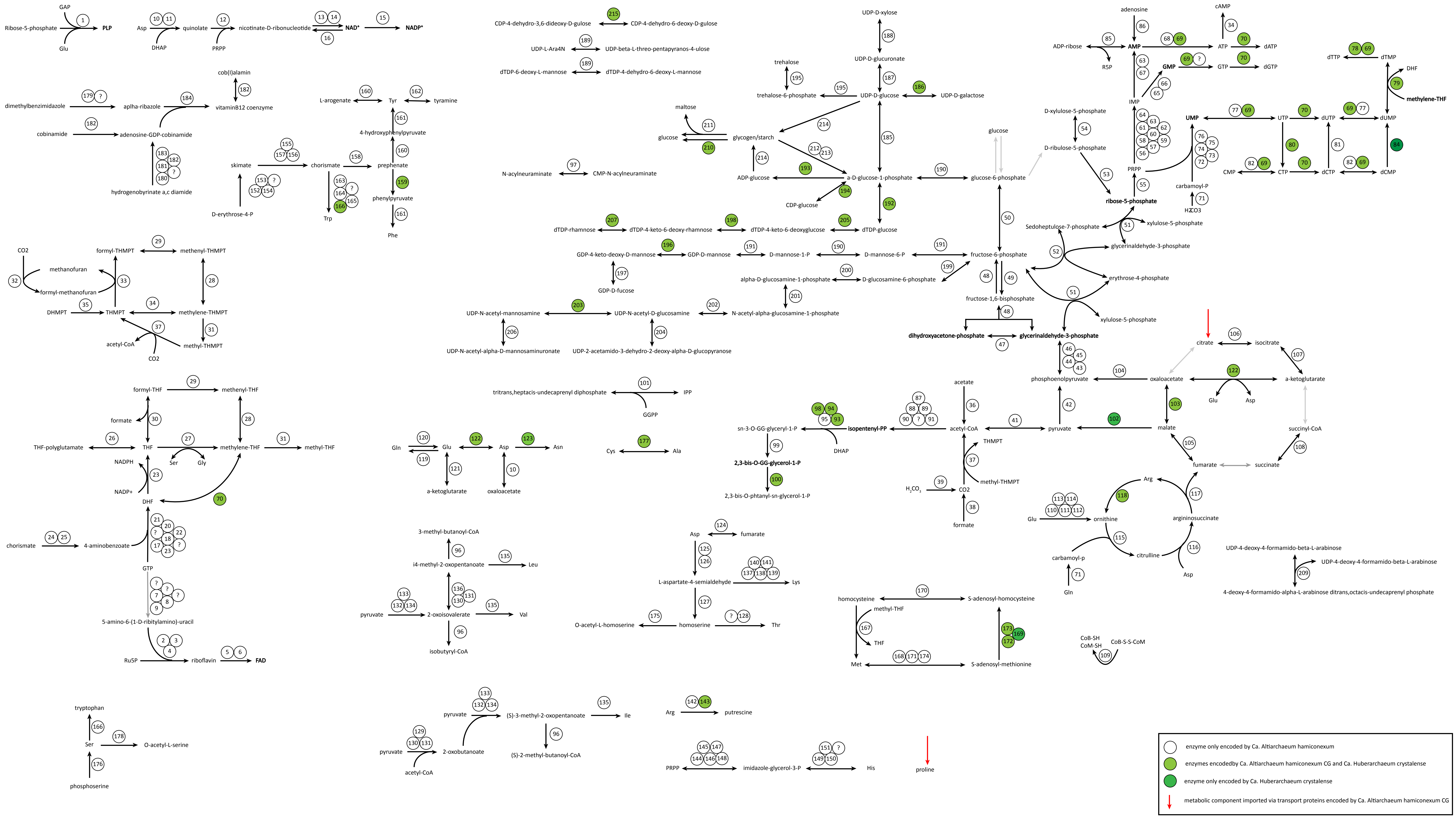
GAP
Ribose-5-phosphate
PLP
Glu
Asp
quinolate
DHAP
PRPP
nicotinate-D-ribonucleotide
NAD+
NADP+
cob(I)alamin
dimethylbenzimidazole
aplha-ribazole
vitaminB12 coenzyme
cobinamide
adenosine-GDP-cobinamide
hydrogenobyrinate a,c diamide
L-arogenate
Tyr
tyramine
4-hydroxyphenylpyruvate
skimate
chorismate
prephenate
phenylpyruvate
D-erythrose-4-P
Trp
Phe
CDP-4-dehydro-3,6-dideoxy-D-gulose
CDP-4-dehydro-6-deoxy-D-gulose
UDP-L-Ara4N
UDP-beta-L-threo-pentapyranos-4-ulose
dTDP-6-deoxy-L-mannose
dTDP-4-dehydro-6-deoxy-L-mannose
N-acylneuraminate
CMP-N-acylneuraminate
UDP-D-xylose
UDP-D-glucuronate
trehalose
trehalose-6-phosphate
UDP-D-glucose
UDP-D-galactose
maltose
glucose
glycogen/starch
ADP-glucose
a-D-glucose-1-phosphate
CDP-glucose
dTDP-glucose
glucose-6-phosphate
dTDP-rhamnose
dTDP-4-keto-6-deoxy-rhamnose
dTDP-4-keto-6-deoxyglucose
GDP-4-keto-deoxy-D-mannose
GDP-D-mannose
D-mannose-1-P
D-mannose-6-P
fructose-6-phosphate
GDP-D-fucose
alpha-D-glucosamine-1-phosphate
D-glucosamine-6-phosphate
fructose-1,6-bisphosphate
UDP-N-acetyl-mannosamine
UDP-N-acetyl-D-glucosamine
N-acetyl-alpha-glucosamine-1-phosphate
UDP-N-acetyl-alpha-D-mannosaminuronate
UDP-2-acetamido-3-dehydro-2-deoxy-alpha-D-glucopyranose
dihydroxyacetone-phosphate
glycerinaldehyde-3-phosphate
adenosine
ADP-ribose
AMP
R5P
ATP
cAMP
dATP
GMP
GTP
dGTP
IMP
D-xylulose-5-phosphate
D-ribulose-5-phosphate
PRPP
ribose-5-phosphate
UMP
UTP
dUTP
dUMP
dTTP
dTMP
DHF
methylene-THF
carbamoyl-P
$H_2CO_3$
CMP
CTP
dCTP
dCMP
Sedoheptulose-7-phosphate
xylulose-5-phosphate
glycerinaldehyde-3-phosphate
erythrose-4-phosphate
xylulose-5-phosphate
CO2
methanofuran
formyl-methanofuran
formyl-THMPT
methenyl-THMPT
DHMPT
THMPT
methylene-THMPT
acetyl-CoA
methyl-THMPT
CO2
tritrans,heptacis-undecaprenyl diphosphate
IPP
GGPP
citrate
isocitrate
a-ketoglutarate
phosphoenolpyruvate
oxaloacetate
Glu
Asp
acetate
sn-3-O-GG-glyceryl-1-P
isopentenyl-PP
acetyl-CoA
pyruvate
malate
succinyl-CoA
DHAP
THMPT
fumarate
succinate
2,3-bis-O-GG-glycerol-1-P
methyl-THMPT
2,3-bis-O-phtanyl-sn-glycerol-1-P
$H_2CO_3$
CO2
formate
Arg
Glu
ornithine
argininosuccinate
citrulline
carbamoyl-p
Asp
Gln
formyl-THF
methenyl-THF
formate
THF-polyglutamate
THF
methylene-THF
methyl-THF
Ser
Gly
NADPH
NADP+
DHF
Gln
Glu
Asp
Asn
Cys
Ala
a-ketoglutarate
oxaloacetate
chorismate
4-aminobenzoate
GTP
3-methyl-butanoyl-CoA
i4-methyl-2-oxopentanoate
Leu
pyruvate
2-oxoisovalerate
Val
isobutyryl-CoA
fumarate
L-aspartate-4-semialdehyde
Lys
homoserine
O-acetyl-L-homoserine
Thr
homocysteine
S-adenosyl-homocysteine
methyl-THF
THF
Met
S-adenosyl-methionine
UDP-4-deoxy-4-formamido-beta-L-arabinose
UDP-4-deoxy-4-formamido-beta-L-arabinose
4-deoxy-4-formamido-alpha-L-arabinose ditrans,octacis-undecaprenyl phosphate
CoB-SH
CoM-SH
CoB-S-S-CoM
5-amino-6-(1-D-ribitylamino)-uracil
Ru5P
riboflavin
FAD
tryptophan
Ser
O-acetyl-L-serine
phosphoserine
pyruvate
(S)-3-methyl-2-oxopentanoate
Ile
(S)-2-methyl-butanoyl-CoA
pyruvate
2-oxobutanoate
acetyl-CoA
Arg
putrescine
PRPP
imidazole-glycerol-3-P
His
proline
enzyme only encoded by Ca. Altiarchaeum hamiconexum
enzymes encodedby Ca. Altiarchaeum hamiconexum CG and Ca. Huberarchaeum crystalense
enzyme only encoded by Ca. Huberarchaeum crystalense
metabolic component imported via transport proteins encoded by Ca. Altiarchaeum hamiconexum CG

**Supplementary Data file 1.**
Table summarizing the genomes used for the maximum likelihood and Bayesian phylogenetic analyses. Includes the NCBI assembly accession ID, taxonomic affiliation as well as CheckM results and number of marker genes identified using phylosift.

**Supplementary Data file 2**
Raw tree file of the maximum likelihood tree displayed in Figure 1A. It includes 186 taxa and was inferred based on an alignment of 34 marker genes with 4224 positions in IQ-tree under the LG+C60+F+R model. Bootstrap support was inferred using an SH-like approximate likelihood ratio test and ultrafast bootstrap support values. Scale bar indicates the average number of substitutions per site.

**Supplementary Data file 3**
Raw tree file of the maximum likelihood analysis based on the recoded alignment used in the analysis shown in Fig. 1. It includes 186 taxa and was inferred based on a SR4-recoded alignment of 34 marker genes with 4224 positions in IQ-tree under the LG+C60+F+R model. Bootstrap support was inferred using an SH-like approximate likelihood ratio test and ultrafast bootstrap support values. Scale bar indicates the average number of substitutions per site.

**Supplementary Data file 4**
Raw tree of the Bayesian phylogenetic analysis based on the concatenated alignment of the 34 marker genes with 4288 sites using 124 representative archaeal taxa. The tree represents a consensus generated from 2 chains, sampling every fifth generation (maxdif: 0.14). Values at branches refer to posterior probability support.

**Supplementary Data file 5**
Raw tree of the Bayesian phylogenetic analysis based on the recoded concatenated alignment of the 34 marker genes with 4288 sites using 124 representative archaeal taxa. The tree represents a consensus generated from 2 chains, sampling every fifth generation (maxdif: 0.14). Values at branches refer to posterior probability support.

### 4. References

1. Darling AE, Jospin G, Lowe E, Matsen IV FA, Bik HM, Eisen JA. PhyloSift: phylogenetic analysis of genomes and metagenomes. *PeerJ* 2014; **2**: e243.
2. Becker EA, Seitzer PM, Tritt A, Larsen D, Krusor M, Yao AI, et al. Phylogenetically driven sequencing of extremely halophilic archaea reveals strategies for static and dynamic osmo-response. *PLoS Genet* 2014; **10**: e1004784.
3. Katoh K, Misawa K, Kuma K, Miyata T. MAFFT: a novel method for rapid multiple sequence alignment based on fast Fourier transform. *Nucleic Acids Res* 2002; **30**: 3059–3066.
4. Probst AJ, Weinmaier T, Raymann K, Perras A, Emerson JB, Rattei T, et al. Biology of a widespread uncultivated archaeon that contributes to carbon fixation in the subsurface. *Nat Commun* 2014; **5**: 5497.
5. Susko E, Roger AJ. On reduced amino acid alphabets for phylogenetic inference. *Mol Biol Evol* 2007; **24**: 2139–2150.
6. Zaremba-Niedzwiedzka K, Caceres EF, Saw JH, Bäckström D, Juzokaite L, Vancaester E, et al. Asgard archaea illuminate the origin of eukaryotic cellular complexity. *Nature* 2017; **541**: 353–358.
7. Guindon S, Dufayard J-F, Lefort V, Anisimova M, Hordijk W, Gascuel O. New algorithms and methods to estimate maximum-likelihood phylogenies: assessing the performance of PhyML 3.0. *Syst Biol* 2010; **59**: 307–321.
8. Minh BQ, Nguyen MAT, von Haeseler A. Ultrafast approximation for phylogenetic bootstrap. *Mol Biol Evol* 2013; **30**: 1188–1195.
9. Lartillot N, Philippe H. A Bayesian mixture model for across-site heterogeneities in the amino-acid replacement process. *Mol Biol Evol* 2004; **21**: 1095–1109.
10. Probst AJ, Ladd B, Jarett JK, Geller-McGrath DE, Sieber CMK, Emerson JB, et al. Differential depth distribution of microbial function and putative symbionts through sediment-hosted aquifers in the deep terrestrial subsurface. *Nat Microbiol* 2018; **3**: 328–336.
11. Edgar RC. Search and clustering orders of magnitude faster than BLAST. *Bioinformatics* 2010; **26**: 2460–2461.
12. Vallenet D, Labarre L, Rouy Z, Barbe V, Bocs S, Cruveiller S, et al. MaGe: a microbial genome annotation system supported by synteny results. *Nucleic Acids Res* 2006; **34**: 53–65.
13. Kanehisa M, Sato Y, Kawashima M, Furumichi M, Tanabe M. KEGG as a reference resource for gene and protein annotation. *Nucleic Acids Res* 2016; **44**: D457–D462.
14. Boeckmann B, Bairoch A, Apweiler R, Blatter M-C, Estreicher A, Gasteiger E, et al. The SWISS-PROT protein knowledgebase and its supplement TrEMBL in 2003. *Nucleic Acids Res* 2003; **31**: 365–370.
15. Probst AJ, Castelle CJ, Singh A, Brown CT, Anantharaman K, Sharon I, et al. Genomic resolution of a cold subsurface aquifer community provides metabolic insights for novel microbes adapted to high $CO_2$ concentrations. *Environ Microbiol* 2017; 19: 459-474.
16. Emerson JB, Thomas BC, Alvarez W, Banfield JF. Metagenomic analysis of a high carbon dioxide subsurface microbial community populated by chemolithoautotrophs and bacteria and archaea from candidate phyla. *Environ Microbiol* 2016; **18**: 1686–1703.
17. Brown CT, Olm MR, Thomas BC, Banfield JF. Measurement of bacterial replication rates in microbial communities. *Nat Biotechnol* 2016; 34: 1256-1263.

18. R Core T. R: A language and environment for statistical computing. *Online Httpwww R-Proj Org* 2016.
19. Ludwig W, Strunk O, Westram R, Richter L, Meier H, Yadhukumar, et al. ARB: a software environment for sequence data. *Nucleic Acids Res* 2004; **32**: 1363–1371.
20. Rudolph C, Wanner G, Huber R. Natural communities of novel archaea and bacteria growing in cold sulfurous springs with a string-of-pearls-like morphology. *Appl Environ Microbiol* 2001; **67**: 2336–2344.
21. Yilmaz LS, Parnerkar S, Noguera DR. mathFISH, a web tool that uses thermodynamics-based mathematical models for in silico evaluation of oligonucleotide probes for fluorescence in situ hybridization. *Appl Environ Microbiol* 2011; **77**: 1118–1122.
